# Subcellular carbohydrate compartmentation and organic acid signatures reveal natural variation in cold acclimation of *Arabidopsis thaliana*

**DOI:** 10.64898/2026.08.31.748218

**Authors:** Vladimir Brodsky, Wolfram Weckwerth, Thomas Nägele

**Affiliations:** LMU München, Faculty of Biology, Plant Evolutionary Cell Biology, Großhaderner Str. 2-4, 82152 Planegg, Germany; University of Vienna, Faculty of Life Sciences, Molecular Systems Biology, Djerassiplatz 1, 1030 Vienna, Austria

**Keywords:** *Arabidopsis thaliana*, natural variation, cold acclimation, leaf development, photosynthesis, subcellular carbon metabolism

## Abstract

Plant cold acclimation emerges from coordinated adjustments in photosynthesis, primary metabolism, and intracellular carbon allocation. Yet, the regulatory role of subcellular metabolite compartmentation in natural variation of cold acclimation remains insufficiently understood. Here, we investigated four *Arabidopsis thaliana* accessions grown either individually or in bulk to determine how growth configuration and genotype shape the metabolism of sugars and organic acids during cold exposure. Using non-aqueous fractionation, we quantified plastidial, cytosolic, and vacuolar sugar pools alongside whole-cell carbohydrates, organic acids, enzyme activities, photosynthetic parameters, and stress markers. A neural-network classifier revealed that subcellular sugar distribution together with sugar amounts and organic acids provided the strongest discriminatory power among accessions, surpassing photosynthetic traits and enzyme activities. Our findings demonstrate that natural variation in cold acclimation is strongly determined by genotype-specific subcellular metabolite architectures, and that the cultivation strategy modulates these intracellular signatures. We conclude that subcellular compartmentation of metabolites represents a cellular control layer for natural variation of cold acclimation and resilience in *Arabidopsis thaliana*.

## Introduction

Plants exhibit remarkable metabolic plasticity, enabling them to buffer environmental fluctuations through the coordinated regulation of photosynthesis, primary metabolism, and intracellular resource allocation (Herrmann et al., 2020; Singh et al., 2020). This plasticity underpins the ability of plants to acclimate to dynamic environmental conditions and has likely been shaped by natural selection across heterogeneous habitats. Consequently, naturally occurring genetic variation provides a valuable framework for identifying the metabolic and physiological mechanisms that determine acclimation capacity. In *Arabidopsis thaliana*, extensive natural variation among accessions originating from distinct habitats has revealed substantial differences in growth, stress tolerance, and metabolic regulation, reflecting adaptation to local environmental conditions (Hannah et al., 2005; Hannah et al., 2006; Verslues and Juenger, 2011; Weigel, 2012).

Central to photosynthetic acclimation is the balance between light-driven energy capture by the photosystems and biochemical carbon fixation through the Calvin– Benson–Bassham cycle (CBBc). The carbohydrates generated by photosynthesis not only constitute the primary products of carbon assimilation but also function as signaling molecules coordinating metabolic acclimation and growth (Schurr et al., 2006; Stitt et al., 2021). Among these, sucrose plays a pivotal role during cold acclimation, acting both as a metabolic hub and a regulator of photosynthetic performance (Strand et al., 2003; Nägele et al., 2012). Sucrose biosynthesis in photosynthetically active leaves is primarily controlled by sucrose phosphate synthase (SPS), after which sucrose can either be exported to sink tissues or hydrolyzed into glucose and fructose by invertases localized in different cellular compartments, including the cytosol, vacuole, plastids, and mitochondria (Xiang et al., 2011). Together with ATP-dependent phosphorylation of glucose and fructose by glucokinases and fructokinases, these reactions establish a cyclic sucrose turnover that has been described both as a metabolic futile cycle (Geigenberger and Stitt, 1991) and as an important mechanism for stabilizing photosynthesis and maintaining metabolic flexibility under fluctuating environmental conditions (Weiszmann et al., 2018).

Under cold temperatures combined with high irradiance, imbalances between absorbed light energy and carbon assimilation increase the risk of reactive oxygen species (ROS) formation and photoinhibition. To mitigate these effects, the photosynthetic apparatus dynamically partitions excitation energy between photochemical conversion, regulated non-photochemical quenching, and non-regulated energy dissipation, thereby balancing photosynthetic efficiency with photoprotection (Roach and Krieger-Liszkay, 2014; Zuo, 2025). Downstream of these photochemical adjustments, metabolic acclimation involves extensive reprogramming of central metabolic pathways, including the metabolism of sugars, organic acids, and amino acids (Hurry et al., 2000; Hoermiller et al., 2022; Saunders et al., 2022). Cold acclimation therefore requires tight coordination between light harvesting, carbon assimilation, sucrose cycling, organic acid metabolism, and intracellular carbon allocation.

While cold-induced metabolic reprogramming has frequently been characterized at the whole-tissue level, increasing evidence indicates that bulk metabolite concentrations provide only a partial representation of cellular metabolic regulation. The distribution of metabolites between plastids, the cytosol, and vacuoles is not merely a passive consequence of cellular organization but represents an additional regulatory layer capable of shaping metabolic fluxes and physiological phenotypes (Hernandez et al., 2023). Compartment-specific metabolite pools may therefore determine how plants coordinate anabolic and catabolic processes, stabilize photosynthesis, buffer redox imbalances, and allocate carbon during environmental fluctuations. This may be particularly relevant for sugars and organic acids, whose cellular functions depend strongly on their localization and exchange between compartments. Consequently, genetically determined differences in intracellular metabolite allocation could contribute to the substantial natural variation in cold acclimation observed among *A. thaliana* accessions.

In addition to genotype and intracellular metabolic organization, the conditions under which plants are cultivated may substantially affect the metabolic state from which acclimation develops. Typically, experimental studies investigating cold acclimation employ highly standardized conditions in which individual plants are cultivated separately to minimize environmental variation and clearly define experimental parameters. Although such designs facilitate mechanistic investigations, their ecological relevance remains uncertain. Natural growth conditions differ not only in soil composition and microclimate but also in the space available per plant and the resulting developmental trajectories. In the field, *A. thaliana* seeds may germinate in densely occupied areas, resulting in competition and mutual effects on photon absorption, nutrient uptake, and water availability. Interactions between developmental stage, plant density, natural genetic variation, and intracellular metabolic regulation therefore remain largely unexplored under standardized laboratory conditions despite their potential relevance for ecological interpretation.

To assess how these factors interact during cold acclimation, the present study investigated four natural *A. thaliana* accessions cultivated either in a *one-plant-per-pot* design or in a *bulk* setup, with several plants growing side by side within the same pot. We specifically examined whether growth configuration and natural genetic variation shape intracellular metabolite allocation during cold exposure and whether such differences are associated with distinct acclimation responses. By combining non-aqueous fractionation with measurements of photosynthetic performance, carbohydrates, organic acids, carbohydrate-metabolizing enzyme activities, and stress and acclimation markers, we characterized cold-induced responses from the whole-cell to the subcellular level. In addition, neural-network-based classification was applied to determine which physiological and metabolic feature groups most effectively discriminate natural accessions and growth configurations. This integrated approach allowed us to test whether compartment-specific metabolite distributions provide discriminatory and mechanistic information beyond conventional whole-cell metabolic and photosynthetic traits and to evaluate the contribution of subcellular carbon allocation to natural variation in cold acclimation.

## Materials and methods

### Plant material and growth conditions

Natural accessions of *Arabidopsis thaliana*, Ct-1 (geographical origin: Catania, Italy; NASC ID: N6674), Fei-0 (geographical origin: St. Maria d. Feiria, Portugal; NASC ID: N22645), Oy-0 (geographical origin: Oystese, Norway; NASC ID: N6824) and Rsch-4 (geographical origin: Rschew/Starize, Russia; NASC ID: N6850), were grown on a 1:1 mixture of GS90 soil and vermiculite in a single plant setup or in a bulk setup. For the bulk growth setup, seeds were randomly sown across the soil and transferred to a greenhouse for 14 days, prior to sampling at midday. Pictures of sampled bulk plants are provided in the supplements (Supplementary Figure SF1). In the single growth setup, plant rosettes were grown isolated, i.e., one plant per pot, in a climate chamber under controlled long-day conditions (8 h/16 h light/dark; 100 μmol m^−2^ s^−1^; 22°C/16°C; 50-60% relative air humidity). After 4 weeks, plants were transferred to long day growth conditions (16 h/8 h light/dark) for additional two weeks before being sampled at midday. Pictures of sampled plants are provided in the supplements (Supplementary Figure SF2). For treatment, plants were then transferred to low temperature and ambient light (LT; 16 h/8 h light/ dark; 125 μmol m^−2^ s^−1^; 4-6°C/4°C, 40-50% relative air humidity), or low temperature and elevated light (LT-EL; 16 h/8 h light/ dark; 250 μmol m^−2^ s^−1^; 6-8°C/4°C, 40-50% relative air humidity). After 7 and 14 days at LT/ LT-EL, single grown plants were sampled at midday. After 10 and 26 days at LT/ LT-EL, bulk plants were sampled at midday. The treatment period for single grown plants, i.e. 7 and 14 days, reflected a typically applied cold acclimation period used by previous studies which resulted in a stable acclimated metabolic homeostasis and a maximal freezing tolerance, see e.g. (Klotke et al., 2004; Distelbarth et al., 2013). For bulk plants, sampling time points were chosen based on induction of inflorescences (after 10 days: bolting stage of first accession Ct-1; after 26 days: bolting stage reached by all accessions). During sampling, plants were immediately quenched in liquid nitrogen and stored at −60°C until further use.

### Net photosynthesis and chlorophyll fluorescence measurements

Net rates of CO_2_ assimilation (*Net PhotoSynthesis*, NPS) and photosynthetic parameters were recorded with increasing light intensities between 0 and 1200 μmol photons m^−2^ s^−1^. Gas exchange and chlorophyll fluorescence were recorded with a WALZ GFS-3000FL system (Heinz Walz GmbH; www.walz.com) coupled to Standard Measuring Head 3010-S with LED Light Source 3041-L. Maximum quantum yield of PSII (F_v_/F_m_) was determined after 15 minutes of dark adaptation of the rosette and subsequently supplying a saturating light pulse. All photosynthetic measurements were conducted at 22 °C and repeated on three independent samples, i.e., single leaf rosettes or bulk grown plants. To estimate physiological relevance of carbon assimilation, NPS rates were temperature-corrected by a factor derived from measurements at low temperature in a previous study (Kitashova et al., 2023).

### Quantification of starch and soluble carbohydrates

Transitory starch and soluble carbohydrate amounts were determined as described before (Brodsky et al., 2025). Dried plant material was suspended in 80% ethanol and incubated at 80°C for 30 minutes. The supernatant, containing soluble sugars, was separated from the starch-containing pellet and transferred to a separate tube for further analysis. Pellets were hydrolyzed in 0.5 N NaOH at 95°C for 60 min and acidified with 1 M CH_3_COOH. The suspension was then subjected to amyloglucosidase digestion. Resulting glucose moieties were quantified photometrically at 540 nm in a coupled glucose-oxidase/peroxidase/dianisidine assay. Ethanol of previously obtained soluble sugar fractions was evaporated, and the dried pellet was solubilized in water. Sucrose was quantified by incubating the samples in 30% KOH at 95°C for 10 minutes before adding 0.14% (w/v) anthrone in 14.6 M H_2_SO_4_ and incubating at 40°C for 30 min. Complexes were photometrically detected and quantified with a calibration curve at 620 nm.

Glucose amounts were obtained in a coupled enzymatic assay of hexokinase and glucose-6-phosphate dehydrogenase (G6PDH), while fructose amounts were quantified via a coupled hexokinase/phosphoglucose isomerase/G6PDH assay. In both cases, NADP^+^ was reduced to yield NADPH+H^+^ which was quantified at 340nm. Starch and soluble carbohydrates were determined in five independent biological replicates.

### Quantification of enzyme activities

Enzyme activities under optimal temperature and substrate saturation (v_max_) were obtained for sucrose phosphate synthase (SPS), glucokinase (GLCK), fructokinase (FRCK), and invertase (INV) as described previously (Kitashova et al., 2023).

Leaf material suspended in 50 mM HEPES pH 7.5, 10 mM MgCl_2_, 1 mM EDTA, 2.5 mM DTT, 10% (v/v) glycerine and 0.1% (v/v) Triton X-100. SPS enzymes were extracted on ice for 20 minutes. Following centrifugation at 4°C with 20,000 g, the supernatant was incubated for 30 min at 25°C with 50 mM HEPES pH 7.5, 15 mM MgCl_2_, 2.5 mM DTT, 35 mM UDP-glucose, 35 mM F6P and 140 mM G6P. The activity assay was stopped by boiling samples with 30% KOH and sucrose amounts were determined with an anthrone assay as described above.

For glucokinase (GLCK) and fructokinase (FRCK) enzyme activity assays, homogenized leaf tissue was extracted in 50 mM Tris pH 8.0, 0.5 mM MgCl_2_, 1 mM EDTA, 1 mM DTT and 1% (v/v) Triton X-100. Samples were centrifuged at 4°C with 20,000 g and the supernatant was combined with a buffer containing 100 mM HEPES pH 7.5, 10 mM MgCl_2_, 2 mM ATP, 1 mM NADP^+^, 0.5 U G6PDH. Activities were determined from changes of NADPH + H^+^ over time after addition of 5 mM glucose or fructose, for respective measurements of v_max:GLCK_ and v_max:FRCK_, at 30°C and 340 nm. Cytosolic (nINV) and vacuolar (aINV) invertase activities were determined in 50 mM HEPES–KOH pH 7.5, 5 mM MgCl_2_, 2 mM EDTA, 1 mM phenylmethylsulfonylfluoride, 1 mM DTT, 10% (v/v) glycerol and 0.1% (v/v) Triton X-100 on ice. The suspension was centrifuged at 4°C with 20,000 g incubated in (I) a nInv-specific reaction buffer with a pH of 7.5, containing 20 mM HEPES–KOH and 100 mM sucrose, or (II) an aInv-specific reaction buffer at ph 4.7, containing 20 mM sodium acetate and 100 mM sucrose. After incubation at 30°C, the reactions were stopped at 95°C. Glucose amounts were obtained in a coupled enzymatic assay of hexokinase and glucose-6-phosphate dehydrogenase (G6PDH). All enzyme activities were determined in three to five independent biological replicates.

To estimate the role of enzyme activities under growth conditions, all measured activities were temperature-corrected applying the Arrhenius equation as described earlier (Kitashova et al., 2023).

### Quantification of carboxylic acids

Absolute amounts of citrate, fumarate and malate were determined using photometric assay kits (Sigma-Aldrich, MAK057, MAK060 and MAK067, www.sigmaaldrich.com). For extraction, double distilled water of 60 °C was added to dried plant material, and the sample was immediately incubated at 95°C for 15 min. After brief centrifugation, the supernatant was used for photometric measurements as described by the manufacturer. Amounts of carboxylic acids were determined in three independent biological replicates.

### Quantification of SOD, MDA and anthocyanins

The activity of superoxide dismutase (SOD) was determined using a photochemical method being based on the inhibition of nitroblue tetrazolium (NBT) reduction. Samples were ground in 1 mL of 100 mM potassium phosphate buffer. The homogenate was incubated on ice for 15 minutes and then centrifuged for 5 minutes at 4°C. For each sample, three replicates and their corresponding blanks were prepared. To prepare the blanks, 12.5 µL of the supernatant was heat-inactivated at 95°C for 10 minutes. Each reaction tube contained 37.5 µL of 100 mM potassium phosphate buffer (or 50 µL water for the water reference), 12.5 µL of sample supernatant (or inactivated blank), 50 µL of 1 mM NBT, 100 µL of 100 mM methionine, and 775 µL of 10 mM potassium phosphate buffer. The reaction was initiated by adding 25 µL of 1 mM riboflavin. Tubes were incubated with open lids for 10 minutes under direct light in a plant cultivation chamber, ensuring consistent light intensity across all samples. Immediately following incubation, absorbance was measured at 560 nm.

For quantification of malondialdehyde (MDA), samples were ground in 1 ml of 80% (v/v) ethanol. The homogenate was incubated on ice for 15 minutes and subsequently centrifuged for 10 minutes at 4°C. A 200 µl aliquot of the supernatant was mixed with 10 µl of 0.2% (w/v) butylated hydroxytoluene (BHT). To sample tubes, 200 µl of TBA+ solution (20% TCA with 0.65% TBA) was added; for the blanks, 200 µl of TBA– solution (20% TCA only) was added instead. The mixtures were incubated at 95°C for 25 minutes, cooled on ice, and then centrifuged for 5 minutes at 4°C. The absorbance of the samples was measured at 440 nm, 532 nm, and 600 nm.

Anthocyanins were extracted from frozen, ground leaf material using 1 ml extraction buffer containing 18% (v/v) 1-propanol and 1% (v/v) HCl in water. Samples were incubated for 2 h at room temperature in darkness and subsequently centrifuged (10 min, 20,000 x g, RT). The supernatant was transferred to cuvettes for spectrophotometric analysis. Absorbance was measured at 537, 650, and 720 nm. Anthocyanin amounts were quantified using a pelargonidine-chloride standard.

### Non-aqueous fractionation

To achieve efficient separation of plant material under different growth conditions, two NAF protocols were applied: a “forward” NAF (fNAF) and a “reverse” NAF (rNAF). Density gradients consisting of mixtures of tetrachloroethylene (TCE) and heptane (7H) were prepared in advance to enable separation of cellular compartments. In this study, gradients of ρ = 1.4, 1.42, 1.47, 1.52, 1.6 and >1.6 g cm^−3^ were used for 7/10D and 14/26D samples. For 0D samples, an additional fraction below 1.4 g cm^−3^ was included (ρ < 1.35, 1.4, 1.45, 1.5, 1.55 and 1.6 g cm^−3^). Samples were either fractionated with increasing (forward, fNAF) or decreasing densities (reverse, rNAF). For fNAF, 6–8 mg of ground, lyophilized plant material (7/10D and 14/26D) was suspended in 1 mL of the lowest-density gradient (e.g., ρ = 1.4 g cm^−3^) in reaction tubes. Samples were sonicated on ice using a Hielscher UP200St-G (Hielscher Ultrasonics ®, www.hielscher.com) sonicator for 10–15 min in bursts of 20–30 s, with 60 s pauses to prevent overheating. The suspension was filtered through a sieve into a new tube and centrifuged at 22,000 × g and 4 °C for 10–15 min. The supernatant, containing compartments corresponding to the applied density, was transferred to a new tube, briefly sonicated (< 20 s), and divided into two equal subfractions. These were dried in a desiccator and subsequently used for enzyme activity assays and soluble sugar quantification. Equal volumes between subfractions were ensured. The remaining pellet was resuspended in the next higher density (e.g., ρ = 1.42 g cm^−3^), briefly sonicated (< 20 s), and centrifuged under the same conditions. This procedure was repeated sequentially up to the highest density (e.g., ρ = 1.6 g cm^−3^).

For rNAF, 6–8 mg of ground, lyophilized plant material (0D) was suspended in the highest-density gradient (e.g., ρ = 1.6 g cm^−3^). Sonication, filtration, and centrifugation were performed as described for fNAF. The supernatant was carefully transferred to a new tube, and heptane was added to adjust the density to the next lower level (e.g., ρ = 1.52 g cm^−3^). After brief sonication (< 20 s) and centrifugation, this stepwise reduction in density was repeated until the lowest fraction (ρ < 1.35 g cm^−3^) was reached. At each step, the pellet contained compartments corresponding to the respective density. The final pellet was resolubilized in 1 ml pure TCE, briefly sonicated (< 20 s), and split into two equal subfractions. These were processed identically to the fNAF subfractions.

For each density fraction, marker enzyme activities and soluble sugars were quantified. Dried samples of the first subfraction were resuspended in 500 μl extraction buffer, briefly sonicated, and incubated on ice for 20 min. After centrifugation (22,000 × g, 4 °C, 10 min), the supernatant was used for enzyme activity assays. Dried samples of the second subfraction were extracted with 650 μl of 80% (v/v) ethanol at 80 °C for 30 min with shaking. Marker enzyme activities were determined for plastids (alkaline pyrophosphatase), cytosol (UDP-glucose pyrophosphorylase) and vacuole (acidic phosphatase) as described previously (Hernandez et al., 2023).

Sucrose content was determined similarly to cellular level measurements with minor modifications. Dried sugar extracts were resuspended in 400 μl ddH_2_O and reacted with 1 ml anthrone reagent. Relative distribution was calculated as percentage contribution of each subfraction.

For hexose quantification, fractions at the last two highest densities were diluted 1:4 with ddH_2_O to avoid saturation. Relative distributions were calculated as percentage contributions of each subfraction.

Subcellular distributions of metabolites were determined using the NAFalyzer app (https://github.com/cellbiomaths/NAFalyzer).

### Statistics and two-layer feedforward network classification

A multi-factorial ANOVA was performed to test the effects of growth type, accession, time point, and condition, including all interaction terms, on experimentally determined variables. Effect sizes were quantified using *partial eta squared*, η^2^, which represents the proportion of variance in the response variable explained by a given effect after accounting for residual (error) variance. Higher η^2^ values indicated a greater contribution of the respective factor or interaction to the observed variation. A summary of the ANOVA results is provided in the supplements (Supplementary Table ST1). Statistical data evaluation was done in R/RStudio (2026.07.1 Build 147), and the *ggplot2* package was used for graphical representation.

A two-layer feedforward network was applied for data classification. The first layer comprised 10 layers of sigmoid hidden neurons, the second layer comprised a *softmax* function for probability estimation. The classification was performed in MATLAB® (R2025b) using the Deep Learning Toolbox.

Classifier performance was evaluated using the area under the receiver operating characteristic curve (AUC), macro-AUC, accuracy, macro-precision, macro-recall, and macro-F1 score. Receiver operating characteristic (ROC) curves were generated for each class by plotting the true-positive rate (sensitivity) against the false-positive rate across classification thresholds, and the AUC was calculated as the area under the resulting ROC curve.

For multiclass classification, ROC curves and AUCs were determined separately for each class using a one-vs-rest approach. The macro-AUC was calculated as the unweighted arithmetic mean of the class-specific AUC values, thereby assigning equal weight to each class.

Accuracy was calculated as the proportion of correctly classified samples among all samples in the independent test set. For each class, precision was calculated as TP/(TP + FP), where TP and FP denote true-positive and false-positive predictions, respectively. The recall was calculated as TP/(TP + FN), where FN denotes false-negative predictions.

The F1 score was calculated as the harmonic mean of precision and recall [2 × (precision × recall)/(precision + recall)]. Macro-precision, macro-recall, and macro-F1 were subsequently calculated as the unweighted arithmetic means of the corresponding class-specific values, ensuring that each class contributed equally to the overall performance measure irrespective of class size. All performance metrics were determined using the independent test dataset (comprising 15% of full data sets). AUC values of 0.5 indicated discrimination at chance level, whereas values approaching 1 indicated increasingly strong discriminatory performance; for accuracy, precision, recall, and F1 score, values approaching 1 indicate better classification performance.

A summary of all data used for statistics and classification is provided in the supplement (Supplementary Table ST2).

## Results

### Cold-induced dynamics of photosynthesis and carbon metabolism are more pronounced in single-grown than in bulk-grown plants

Under ambient growth conditions, all tested accessions showed a similar photosynthetic performance. However, during cold exposure, photosynthetic parameters revealed significant differences between single-grown and bulk-grown *Arabidopsis* accessions. Maximum quantum yield of photosystem II (Fv/Fm), effective quantum yield of PSII (Y(II)), as well as regulated (Y(NPQ)) and unregulated (Y(NO)) non-photochemical quenching significantly differed between both growth types and between conditions (ANOVA, p < 0.001; Supplementary Table ST1). Significant accession-specific differences were observed for Y(II), Y(NPQ) and Y(NO) whereas Fv/Fm did not differ between accessions (ANOVA, p > 0.05). As a trend, Fv/Fm, Y(II) and Y(NPQ) were found to be lower under cold/elevated light than under cold/ambient light (Fig. 1 A-C) while Y(NO) showed an inverse trend (Fig. 1 D).

**Figure 1.**
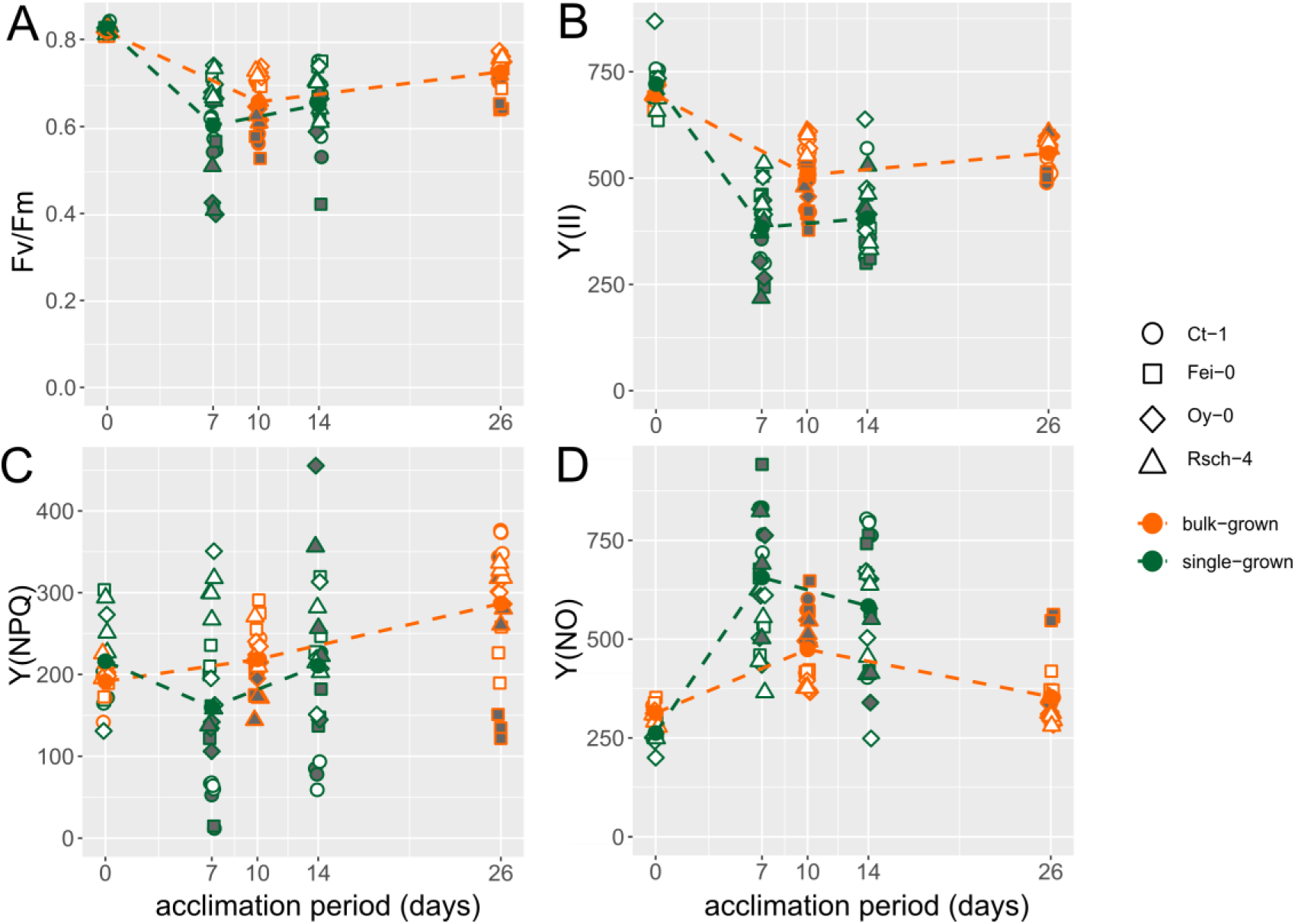
Dynamics of photosynthetic acclimation in natural accessions of *Arabidopsis thaliana* during cold acclimation. Symbols filled white: ambient light (125 µE). Symbols filled grey: elevated light (250 µE). Orange: bulk-grown plants; green: single-grown plants. Symbols represent accessions. Circles: Ct-1; squares: Fei-0; diamonds: Oy-0; triangles: Rsch-4. ANOVA results are provided in the supplements (Supplementary Table ST1).

However, although a significant effect was observed between different growth types, and cold-induced effects were more pronounced in single-grown than in bulk-grown plants, the dynamics were similar between bulk-grown and single-grown plants. Both Fv/Fm and Y(II) decreased until 7 days at 4°C before both parameters were stabilized until 14 days and 26 days in single-grown and bulk-grown plants, respectively (Fig. 1 A, B). Also, for Y(NO) both growth types showed similar dynamics with an initial increase followed by a subsequent decrease until the late acclimation phase (Fig. 1 D). Only dynamics of Y(NPQ) differed between both growth types, showing a constant increase in bulk-grown plants and a decrease followed by an increase in single-grown plants (Fig. 1 C).

Fresh weight-to-dry weight (FW) ratios showed a pronounced temporal decline, with the highest ratios and greatest variability observed at the initial time point, followed by markedly lower values at subsequent measurements (Fig. 2 A). The decline was particularly pronounced between the initial measurement and 7 days, after which FW ratios remained comparatively low. Differences between growth types were most apparent at the initial time point and became less pronounced at later measurements. Net photosynthesis (NPS) also varied strongly across time points and between growth types (Fig. 2 B). Cold-induced dynamics differed between growth types, as indicated by a strong growth type x time point interaction (ANOVA, p < 0.001, partial η² = 0.498). In contrast, the growth type x condition interaction and all remaining interactions involving accessions were not significant (p ≥ 0.05).

**Figure 2.**
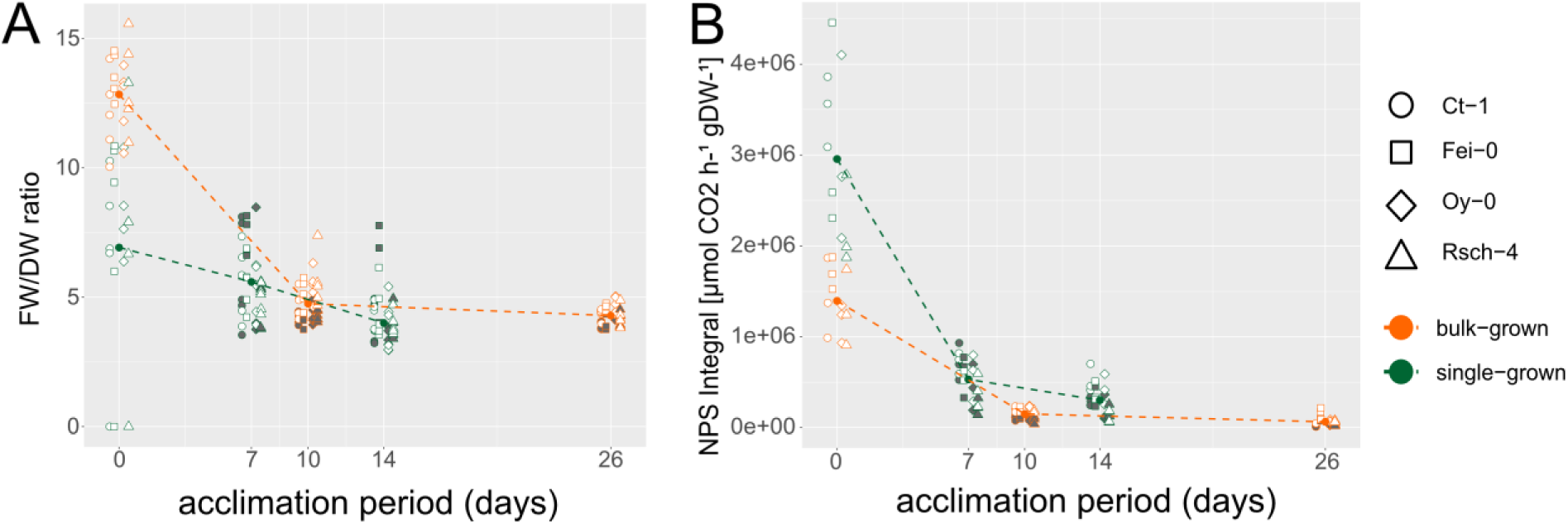
Fresh weight to dry weight ratio (A) and net photosynthesis (B) of leaf tissue across time points and growth types. NPS values represent integrals of CO_2_ assimilation rates over recorded light curves. Symbols filled white: ambient light (125 µE). Symbols filled grey: elevated light (250 µE). Orange: bulk-grown plants; green: single-grown plants. Symbols represent accessions. Circles: Ct-1; squares: Fei-0; diamonds: Oy-0; triangles: Rsch-4.

The quantification of central carbohydrates revealed significant effects of the growth type on accumulation dynamics during cold exposure. Starch accumulated linearly during cold exposure of single grown plants while bulk-grown plants showed a significant increase during the initial 10 days and a subsequent plateau-like dynamic until 26 days (Figure 3 A). Similarly, also sucrose amounts showed a linear increase over the initial 14 days of cold acclimation in single-grown plants, whereas amounts in bulk-grown plants remained constant between 10 and 26 days of cold exposure (Figure 3 B).

**Figure 3.**
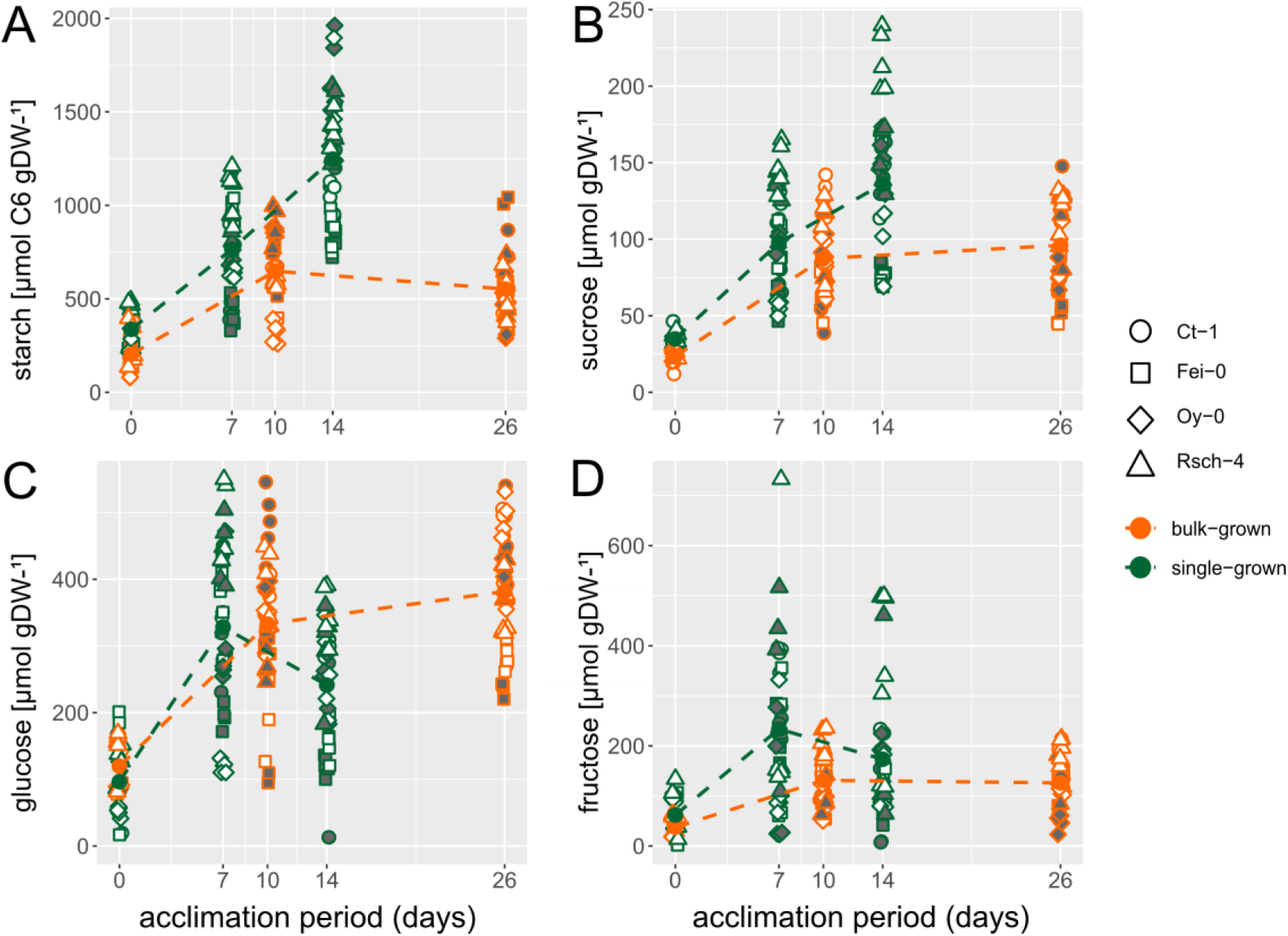
Cold-induced carbohydrate dynamics in natural accessions of *Arabidopsis thaliana* during cold acclimation. Symbols filled white: ambient light (125 µE). Symbols filled grey: elevated light (250 µE). Orange: bulk-grown plants; green: single-grown plants. Symbols represent accessions. Circles: Ct-1; squares: Fei-0; diamonds: Oy-0; triangles: Rsch-4.

Both free hexoses glucose and fructose showed similar amounts and dynamics in single-grown and bulk-grown plants, with an initial cold-induced increase and a constant amount until 14 and 26 days, respectively (Figure 3 C, D). Interestingly, only for starch a significant effect was observed between ambient and elevated light treatments with significantly higher amounts in elevated light samples (ANOVA p < 0.001, Supplementary Table ST1). In summary, this suggested that both analysed growth types could be differentiated by the accumulation dynamics of starch and sucrose, whereas light intensity at low temperature was most significantly reflected by starch accumulation.

Next, dynamics of organic acids citrate, malate and fumarate were quantified which, in addition to carbohydrates, represent another central metabolic carbon pool. Single-grown plants accumulated significantly higher amounts of organic acids than bulk-grown plants across all time points, also before cold acclimation (Figure 4). In bulk-grown plants, citrate and malate showed a continuous accumulation during cold exposure, while fumarate declined. In single-grown plants, citrate amounts were found to peak after 7 days at 4°C followed by a decline until 14 days of cold exposure (Figure 4 A). While malate amounts continuously declined, fumarate amounts significantly increased during 14 days of cold exposure (Figure 4 B, C).

**Figure 4.**
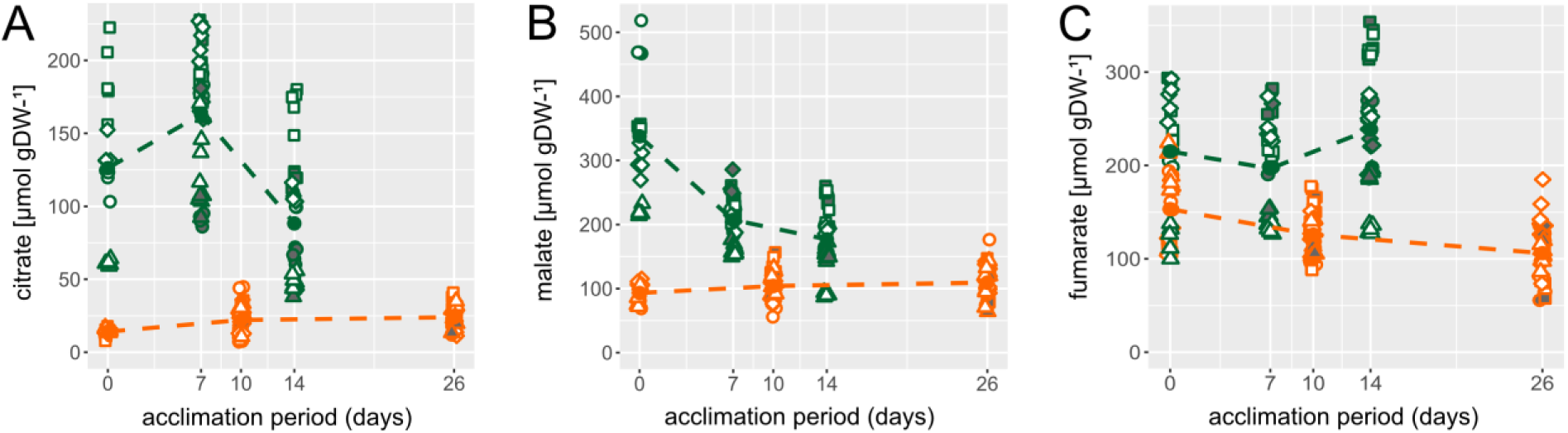
Cold-induced dynamics of organic acids in natural accessions of *Arabidopsis thaliana* during cold acclimation. Symbols filled white: ambient light (125 µE). Symbols filled grey: elevated light (250 µE). Orange: bulk-grown plants; green: single-grown plants. Symbols represent accessions. Circles: Ct-1; squares: Fei-0; diamonds: Oy-0; triangles: Rsch-4.

### Metabolic stress and acclimation markers suggest growth-type specific cold acclimation trajectories

To estimate cold-induced stress levels and acclimation output, the activity of the enzyme superoxide dismutase (SOD) was quantified together with amounts of malondialdehyde (MDA) and anthocyanins. The stress marker SOD constantly increased during cold exposure in both growth types while activities were constantly higher in single-grown plants (Figure 5 A). Similarly, also MDA levels increased in both growth types but reached higher levels in single-grown plants while a plateau was reached after 10 days in bulk-grown plants (Figure 5 B). Similar dynamics were observed for anthocyanins which showed saturation-like dynamics on bulk-grown plants while accumulation became rather linear or exponential in single-grown plants until 14 days of cold exposure (Figure 5 C). Growth light had a significant impact on the MDA and anthocyanin levels, resulting in higher amounts of both compounds under elevated light compared to ambient light (Figure 5 B, C).

**Figure 5.**
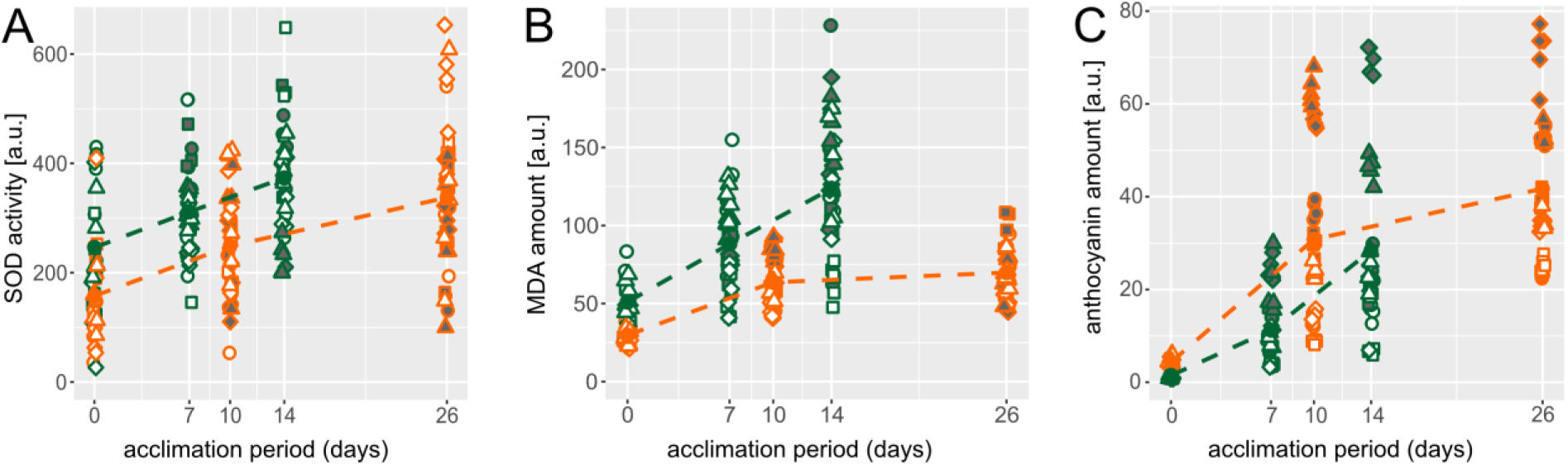
Cold-induced dynamics of stress markers in natural accessions of *Arabidopsis thaliana* during cold acclimation. Symbols filled white: ambient light (125 µE). Symbols filled grey: elevated light (250 µE). Orange: bulk-grown plants; green: single-grown plants. Symbols represent accessions. Circles: Ct-1; squares: Fei-0; diamonds: Oy-0; triangles: Rsch-4.

### Accession × Growth Type Interaction in Subcellular Metabolic Regulation

Previous studies have shown that cold exposure and acclimation are associated with a significant shift of carbohydrates between subcellular compartments (for an overview, see e.g. (Pommerrenig et al., 2018)). To reveal whether cold-induced subcellular carbohydrate allocation was affected by the growth type, a non-aqueous fractionation protocol was applied to separate plastidial from cytosolic and vacuolar carbohydrates (Hernandez et al., 2023). Sugar allocation between the three compartments was found to significantly differ between single-grown and bulk-grown plants (Figure 6). Most pronounced effects were observed for plastidial sucrose allocation (Figure 6 A). In single-grown plants, sucrose accumulated significantly in plastids until 7 days and remained constant until 14 days at 4°C (Figure 6 A). In contrast, bulk-grown plants were found to be depleted in plastidial sucrose until 10 days at 4°C before recovering to pre-acclimation levels. Opposing trends were observed for cytosolic sucrose which indicated a growth-type dependent differential sucrose allocation between both compartments (Figure 6 A, D).

**Figure 6.**
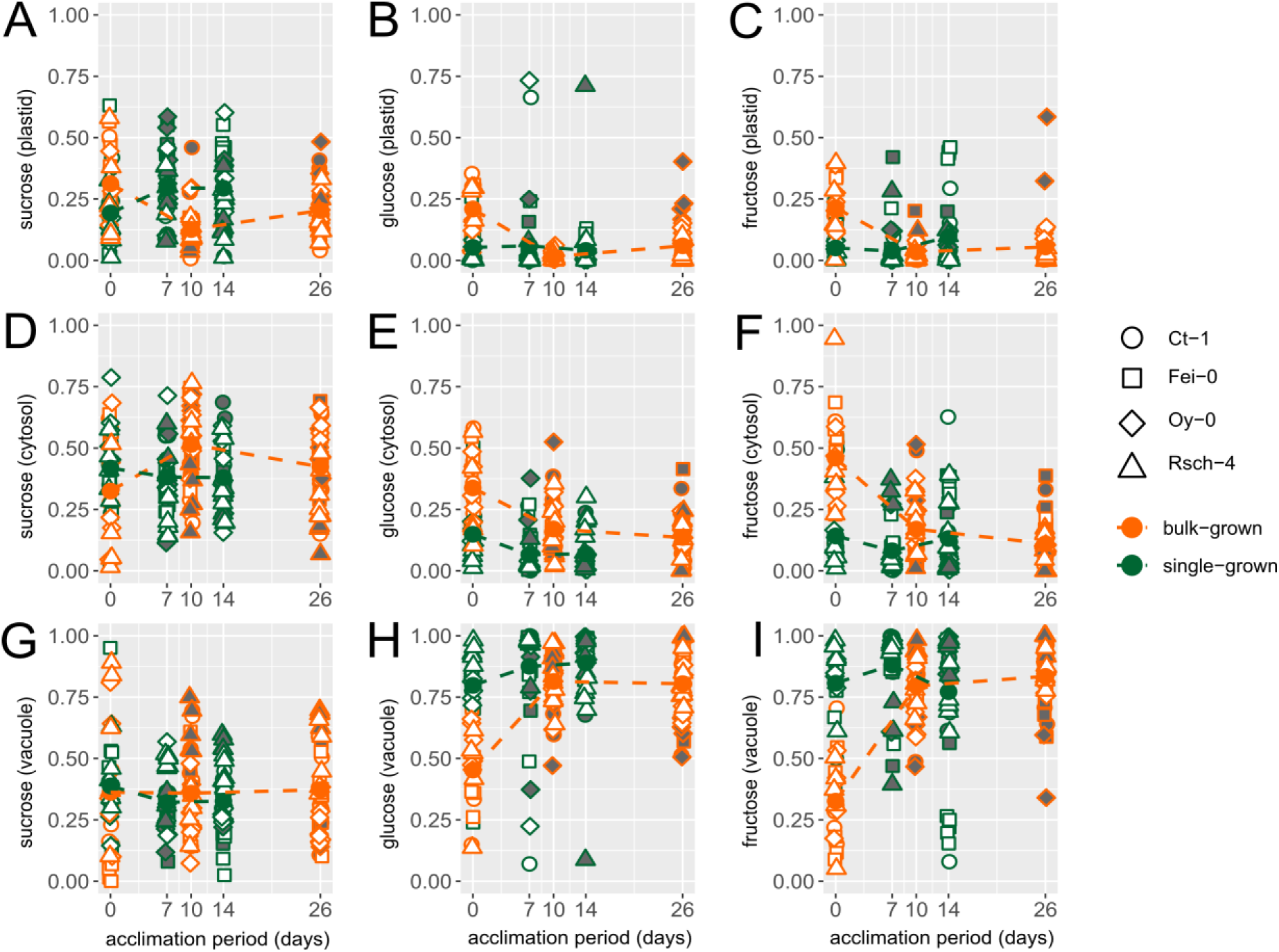
Subcellular sugar allocation during cold acclimation in natural accessions of Arabidopsis thaliana during cold acclimation. Symbols filled white: ambient light (125 µE). Symbols filled grey: elevated light (250 µE). Orange: bulk-grown plants; green: single-grown plants. Symbols represent accessions. Circles: Ct-1; squares: Fei-0; diamonds: Oy-0; triangles: Rsch-4.

Also, dynamics of hexose allocation differed significantly between both growth types. While single-grown plants showed a slight increase of vacuolar hexoses under low temperature, this increase was more pronounced in bulk-grown plants which almost doubled their vacuolar hexose proportions during the first 10 days of cold exposure (Figure 6 H, I). In addition to the depletion of cytosolic hexoses, which was also observed for single-grown plants, bulk-grown plants also shifted hexoses from plastids into the vacuole (Fig 6 B, C). In summary, dynamics of plastidial sucrose allocation differed significantly between growth-types while cold-induced hexose allocation to the vacuole was found to be more pronounced in bulk-grown plants.

A Pearson correlation analysis revealed significantly different regulation of the central carbohydrate metabolism in single and bulk-grown plants (Figure 7). While total amounts of starch and sugars significantly correlated with metabolic enzyme activities (sucrose phosphate synthase, invertases, hexokinases) in plants of both growth types, subcellular sugar distribution was found to correlate more significantly with enzyme activities in bulk-grown than in single-grown plants. Both FrcK and GlcK activities significantly correlated with cytosolic and plastidial fructose and glucose amounts in bulk-grown plants (Figure 7 B) while this effect was weaker, or not significant at all, in single-grown plants (Figure 7 A). Further, while SPS activity negatively correlated with plastidial sucrose in single-grown plants, a positive correlation was observed in bulk-grown plants. Activities of sucrose-hydrolyzing invertases positively correlated with plastidial and cytosolic hexoses while a negative correlation was observed for vacuolar hexoses in bulk-grown plants (Figure 7 B). In single-grown plants, vacuolar sucrose correlated positively with acidic invertase activities (Figure 7 A).

**Figure 7.**
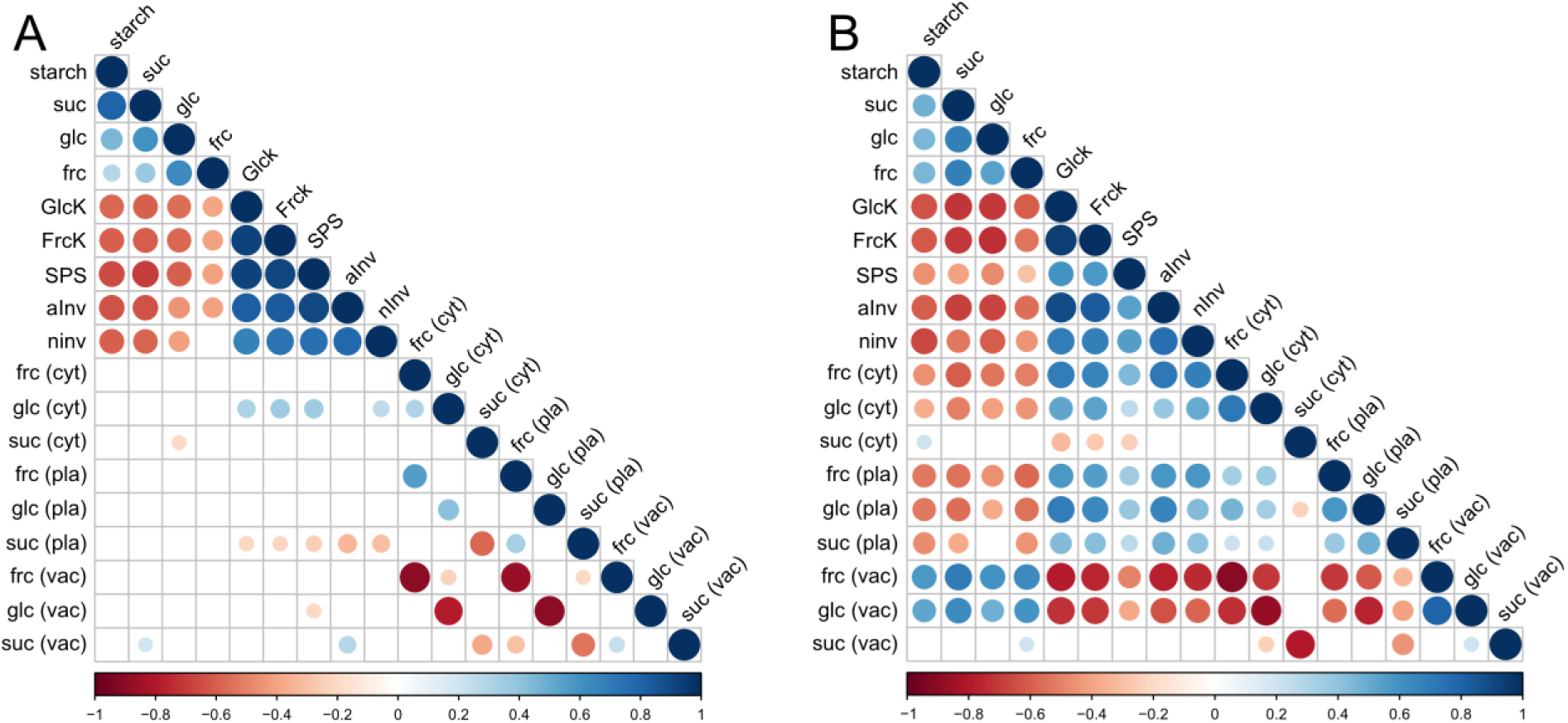
Correlation analysis of sugar metabolism in single-grown (A) and bulk-grown (B) *Arabidopsis* accessions during cold acclimation. Colours indicate positive (blue) or negative (red) Pearson correlation coefficients as shown by the colour bars on the bottom of each plot. Blank fields: non-significant correlation (p > 0.05).

In bulk-grown plants, total sugar amounts showed a more significant correlation with subcellular sugar distributions than in single-grown plants. Plastidial and cytosolic sugar proportions were negatively correlated with total sugar amounts while the vacuolar proportions were positively correlated (Figure 7 B). Also, the correlation between plastidial and cytosolic sugar distribution was more significant in bulk-grown than in single-grown plants. Plastidial sugar proportions were positively correlated with the cytosol while they negatively correlated with the vacuolar proportions. In summary, this suggested that subcellular sugar allocation was differentially regulated in both growth types, and particularly plastidial sucrose allocation showed a significantly reciprocal trend.

### Central enzyme activities of carbohydrate metabolism show growth type-dependent patterns during cold exposure

Enzymatic rates of carbohydrate interconversion were quantified under substrate saturation revealing estimates of v_max_ values of the enzymes sucrose phosphate synthase (SPS), neutral and acidic invertase (aInv, nInv), glucokinase (GlcK) and fructokinase (FrcK), respectively. All enzyme activities were determined at optimal assay temperature and subsequently adjusted to the respective growth temperature using the Arrhenius equation (Kitashova et al., 2023). This resulted in a significant (temperature-induced) drop of all adjusted activities during cold exposure (ANOVA, p < 0.001, Figure 8). Across all tested enzyme activities, the most significant growth-type effect was observed for SPS activities (Figure 8 A). Bulk-grown plants showed significantly lower SPS activities than single-grown plants across all tested time points, i.e., before and during cold exposure (ANOVA, p < 0.001). Except for GlcK, also all other quantified enzyme activities differed significantly between both growth types, but the effect was weaker compared to SPS (ANOVA, p < 0.05, Supplementary Table ST1).

**Figure 8.**
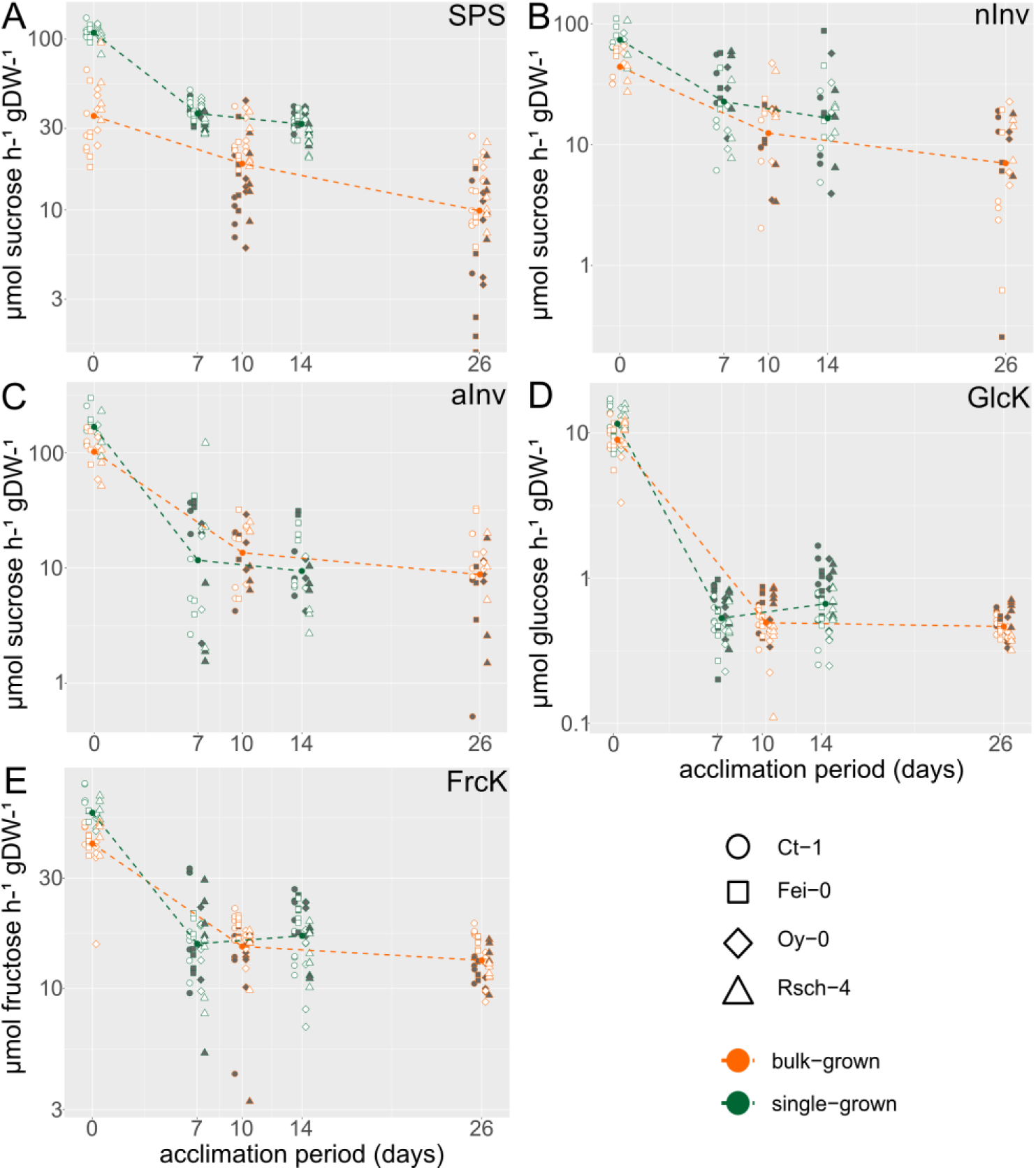
Dynamics of enzyme activities during cold acclimation. **(A)** Sucrose phosphate synthase activities, **(B)** neutral invertase activities, **(C)** acidic invertase activities, **(D)** glucokinase activities, **(E)** fructokinase activities. Symbols filled white: ambient light (125 µE). Symbols filled grey: elevated light (250 µE). Orange: bulk-grown plants; green: single-grown plants. Symbols represent accessions. Circles: Ct-1; squares: Fei-0; diamonds: Oy-0; triangles: Rsch-4.

**Figure 9.**
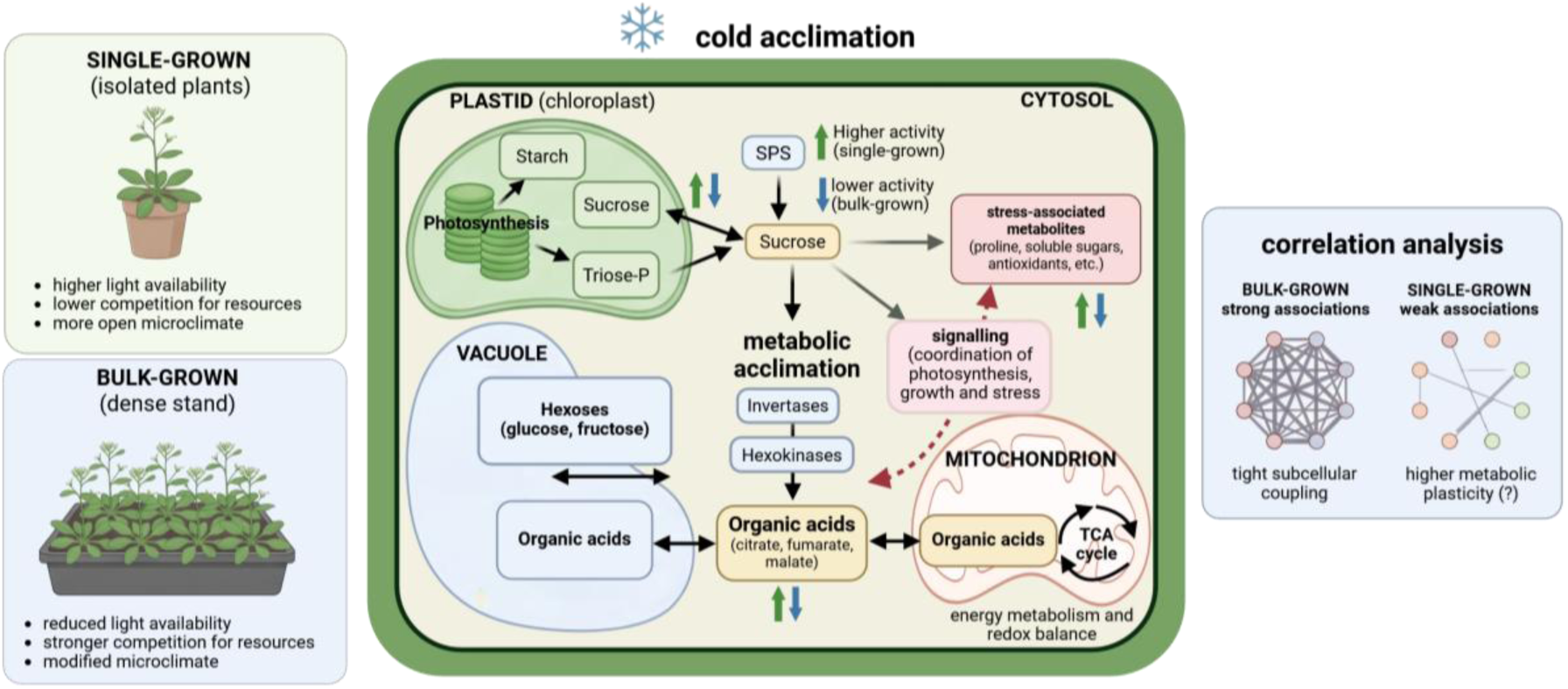
Accessions x growth setup modulates cold acclimation by reshaping metabolic responses and intracellular carbon allocation in *Arabidopsis*. Single-grown and bulk-grown plants experience distinct growth environments that influence photosynthesis, sucrose metabolism, stress-associated metabolites, organic acid metabolism, and subcellular carbohydrate allocation. Arrows indicate metabolic or regulatory relationships, while green and blue arrows denote comparatively stronger responses in single-and bulk-grown plants, respectively. Correlation analysis indicates stronger associations between enzyme activities and subcellular metabolite pools in bulk-grown plants, consistent with tighter metabolic coupling, whereas weaker associations in single-grown plants may indicate greater metabolic plasticity. Created in BioRender. Naegele, T. (2026) https://BioRender.com/g6v8795

### A neural net-based classification reveals organic acids as metabolic biomarkers for natural variation and growth-types during of cold acclimation

To reveal the conservation of cold-induced dynamics of photosynthesis and metabolism across growth setups and accessions, a classifier analysis was performed using a two-layer feedforward deep neural net classifier comprising sigmoid transfer functions in a hidden layer to recognize nonlinearities. The dataset was split in variable categories (i) photosynthesis, (ii) enzymes, (iii) carbohydrates, (iv) organic acids, (v) stress and acclimation markers, and (vi) subcellular sugar distribution. The classifier was trained using 70% of the data, before it was validated and tested each with 15% of the data. To compare the performance of the classifier on the data sets, the area under the curve (AUC) was calculated for each Receiver Operating Characteristic (ROC) curve (Tables I and II). The macro-AUC was calculated as the mean value of all single class AUCs for each classification problem, i.e., “accessions” (Table I) and “growth-type” (Table II).

**Table I.** Classification performance for discrimination between accessions based on different feature groups. Class-specific areas under the receiver operating characteristic curve (AUCs) are reported for each Arabidopsis accession (Ct-1, Fei-0, Oy-0, and Rsch-4) using a one-vs-rest classification approach. The macro-AUC represents the unweighted mean of the four accession-specific AUC values and provides an overall measure of the discriminatory performance of each feature group. AUC values of 0.5 indicate performance at chance level, whereas values approaching 1.0 indicate increasingly strong discrimination. The classifier based on all measured features is shown as a reference for comparison with classifiers based on individual feature groups. A detailed summary of classifier performance metrics is provided in the supplements (Supplementary Table ST3).

| feature group | accession | AUC | macro-AUC |
| --- | --- | --- | --- |
| all features | Ct-1 | 0.463 | 0.485 |
|  | Fei-0 | 0.577 |  |
|  | Oy-0 | 0.447 |  |
|  | Rsch-4 | 0.451 |  |
| photosynthesis | Ct-1 | 0.407 | 0.423 |
|  | Fei-0 | 0.469 |  |
|  | Oy-0 | 0.398 |  |
|  | Rsch-4 | 0.416 |  |
| enzymes | Ct-1 | 0.357 | 0.432 |
|  | Fei-0 | 0.478 |  |
|  | Oy-0 | 0.543 |  |
|  | Rsch-4 | 0.348 |  |
| sugars | Ct-1 | 0.708 | 0.781 |
|  | Fei-0 | 0.824 |  |
|  | Oy-0 | 0.812 |  |
|  | Rsch-4 | 0.780 |  |
| **organic acids** | Ct-1 | 0.925 | 0.841 |
|  | Fei-0 | 0.815 |  |
|  | Oy-0 | 0.667 |  |
|  | Rsch-4 | 0.958 |  |
| **stress and acclimation markers** | Ct-1 | 0.511 | 0.612 |
|  | Fei-0 | 0.749 |  |
|  | Oy-0 | 0.646 |  |
|  | Rsch-4 | 0.542 |  |
| **subcellular sugar distribution** | Ct-1 | 0.665 | 0.729 |
|  | Fei-0 | 0.715 |  |
|  | Oy-0 | 0.680 |  |
|  | Rsch-4 | 0.857 |  |

**Table II.** Classification performance for discrimination between single and bulk growth types based on different feature groups. Class-specific areas under the receiver operating characteristic curve (AUCs) are reported for single and bulk growth using an independent test set (15% of full data sets). The macro-AUC represents the unweighted mean of the AUC values obtained for the two growth types and provides an overall measure of the discriminatory performance of each feature group. An AUC of 0.5 indicates discrimination at chance level, whereas values approaching 1.0 indicate increasingly strong discriminatory performance. The classifier incorporating all measured features is included as a reference for comparison with classifiers based on individual feature groups. A detailed summary of classifier performance metrics is provided in the supplements (Supplementary Table ST4).

| feature group | growth-type | AUC | macro-AUC |
| --- | --- | --- | --- |
| all features | single | 0.769 | 0.620 |
|  | bulk | 0.471 |  |
| photosynthesis | single | 0.500 | 0.563 |
|  | bulk | 0.625 |  |
| enzymes | single | 0.757 | 0.699 |
|  | bulk | 0.641 |  |
| sugars | single | 0.830 | 0.829 |
|  | bulk | 0.828 |  |
| organic acids | single | 1 | 1 |
|  | bulk | 1 |  |
| stress and acclimation markers | single | 0.964 | 0.971 |
|  | bulk | 0.978 |  |
| subcellular sugar distribution | single | 0.835 | 0.830 |
|  | bulk | 0.826 |  |

The discriminatory performance the classifier varied substantially depending on the feature group used for accession classification (Table I). Classification performance was assessed for the accessions Ct-1, Fei-0, Oy-0, and Rsch-4 using class-specific one-vs-rest receiver operating characteristic (ROC) analyses. The classifier incorporating all measured features showed limited discriminatory performance, with a macro-AUC of 0.485. The accession-specific AUCs ranged from 0.447 for Oy-0 to 0.577 for Fei-0, while Ct-1 and Rsch-4 yielded AUCs of 0.463 and 0.451, respectively. Thus, combining all available features did not result in effective discrimination among the four accessions. Similarly, classifiers based exclusively on photosynthetic parameters or enzyme-related features showed low overall discriminatory performance. Photosynthetic features resulted in the lowest macro-AUC among the investigated feature groups (0.423), with class-specific AUCs ranging from 0.398 for Oy-0 to 0.469 for Fei-0. The enzyme-based classifier showed a comparable macro-AUC of 0.432. Within this feature group, Oy-0 showed the highest AUC (0.543), whereas Ct-1 and Rsch-4 showed comparatively low AUCs of 0.357 and 0.348, respectively. In contrast, substantially higher classification performance was obtained when sugar-related variables were used as input features. The sugar-based classifier achieved a macro-AUC of 0.781, with consistently elevated class-specific AUCs across all accessions. AUC values ranged from 0.708 for Ct-1 to 0.824 for Fei-0, with values of 0.812 and 0.780 obtained for Oy-0 and Rsch-4, respectively. Subcellular sugar distribution also showed considerable discriminatory capacity, resulting in a macro-AUC of 0.729. This feature group was particularly effective for the classification of Rsch-4, which exhibited an AUC of 0.857, whereas AUCs for Ct-1, Fei-0, and Oy-0 ranged between 0.665 and 0.715. The highest overall classification performance was obtained using organic acids. The corresponding classifier achieved a macro-AUC of 0.841, exceeding both the classifier based on all features and all other individual feature groups. Particularly high class-specific AUCs were observed for Rsch-4 (0.958) and Ct-1 (0.925), followed by Fei-0 (0.815). Classification of Oy-0 was less pronounced but remained above chance level, with an AUC of 0.667. Thus, the discriminatory information provided by organic acids differed among accessions but was particularly strong for Ct-1 and Rsch-4. Stress and acclimation markers showed intermediate discriminatory performance, with a macro-AUC of 0.612. Among the four accessions, Fei-0 exhibited the highest AUC (0.749), followed by Oy-0 (0.646), whereas Ct-1 (0.511) and Rsch-4 (0.542) showed values close to chance-level discrimination. Taken together, classification performance showed a pronounced dependence on the type of input features. Organic acids provided the highest overall discriminatory performance (macro-AUC = 0.841), followed by sugars (0.781) and subcellular sugar distribution (0.729). In contrast, the combined feature set (0.485), enzyme-related features (0.432), and photosynthetic parameters (0.423) showed substantially lower macro-AUCs.

The ability of the classifier to discriminate between single and bulk growth types varied markedly depending on the feature group used for classification (Table II). The classifier incorporating all measured features showed moderate overall discriminatory performance, with a macro-AUC of 0.620. However, classification performance differed substantially between the two growth types. Single growth was discriminated against with an AUC of 0.769, whereas the AUC for bulk growth was 0.471 and therefore close to chance level. Thus, the combined feature set provided considerably greater discriminatory information for single than for bulk growth within the independent test set. Photosynthetic parameters alone showed similarly limited overall performance, resulting in a macro-AUC of 0.563. The AUC for single growth was 0.500, corresponding to chance-level discrimination, whereas bulk growth showed a moderately higher AUC of 0.625. Enzyme-related features provided greater discriminatory capacity, with a macro-AUC of 0.699. In this case, single growth showed an AUC of 0.757, compared with 0.641 for bulk growth. A substantial increase in classification performance was observed when sugar-related variables were used as input features. The sugar-based classifier achieved a macro-AUC of 0.829, with highly similar class-specific AUC values for single (0.830) and bulk (0.828) growth. This balanced performance indicates that sugar-related features contained considerable discriminatory information for both growth types rather than predominantly distinguishing only one of the two classes. A similarly high and balanced classification performance was obtained from subcellular sugar distribution, with an AUC of 0.835 for single growth and a macro-AUC of 0.830. The strongest discriminatory performance was observed for organic acids and stress and acclimation markers. Organic acid features resulted in perfect separation of the test samples, with AUC values of 1.000 for both single and bulk growth and, consequently, a macro-AUC of 1.000. Stress and acclimation markers also showed near-perfect discriminatory performance, achieving AUC values of 0.964 and 0.978 for single and bulk growth, respectively, and a macro-AUC of 0.971. These feature groups therefore substantially outperformed the classifier incorporating all measured features. Overall, the results revealed pronounced differences in the discriminatory information associated with the investigated feature groups. Organic acids provided the highest classification performance (macro-AUC = 1.000), followed by stress and acclimation markers (0.971), subcellular sugar distribution (0.830), and sugars (0.829). Enzyme-related features showed intermediate performance (0.699), whereas the complete feature set (0.620) and photosynthetic parameters (0.563) exhibited comparatively limited discrimination.

## Discussion

Plant cold acclimation represents a tightly coordinated physiological process that integrates photochemical adjustment, primary metabolism, and intracellular carbon allocation to maintain photosynthetic performance under low, and even freezing, temperatures (Xin and Browse, 2000). While these processes have traditionally been investigated under highly standardized laboratory conditions, the present study demonstrates that the experimental growth setup itself substantially influences the observed acclimation phenotype. Importantly, the differences between single-grown and bulk-grown *Arabidopsis thaliana* plants were not restricted to individual physiological parameters but extended across photosynthesis, central carbon metabolism, enzyme activities, stress markers, and subcellular metabolite compartmentation. Together, these observations suggest that growth conditions modify the regulatory organization of acclimation rather than merely affecting the magnitude of individual metabolic responses.

Although the quantitative responses differed considerably between growth setups, both cultivation strategies followed a similar temporal sequence of acclimation. Photosystem II efficiency initially declined before stabilizing during prolonged cold exposure, while carbohydrates and stress-associated metabolites accumulated over time. Such conservation of the overall response trajectory indicates that the fundamental acclimation programme remains robust across growth environments, maybe due to a conserved underlying perception and signalling network (Ding et al., 2020). In contrast, the consistently stronger responses observed in single-grown plants indicate that the physiological operating point from which acclimation is initiated differs between growth setups. Rather than activating distinct acclimation mechanisms, the two cultivation strategies therefore appear to regulate the same underlying network at different intensities. This might imply that standardized growth conditions influence the quantitative interpretation of acclimation without necessarily altering its (qualitative) regulatory network.

### Growth conditions shape sucrose metabolism during cold acclimation

Significant differences between both growth types during cold acclimation were observed for sucrose metabolism. Continuous sucrose accumulation together with consistently higher sucrose phosphate synthase activities characterized single-grown plants, whereas bulk-grown plants reached an apparent metabolic steady state after the initial acclimation phase. Sucrose plays a central role during cold acclimation because it simultaneously functions as the principal transport carbohydrate, an osmoprotectant, and an important signalling metabolite coordinating photosynthesis and growth (Savitch et al., 2000; Strand et al., 2000). Although it cannot be excluded that the contrasting accumulation dynamics likely reflect differences in carbon allocation strategies, also, lowered photosynthetic carbon assimilation rates in bulk grown plants might be a reason (see Figure 2 B). The higher SPS capacity observed in isolated plants provides a plausible mechanistic basis for sustained sucrose accumulation, whereas the plateau observed in bulk-grown plants suggests that carbon export, utilization, and/or intracellular redistribution becomes increasingly balanced with carbon assimilation during prolonged acclimation.

### Growth setup shapes subcellular carbohydrate partitioning and metabolic coordination

An important conceptual advance of the present study is the observation that growth setup substantially alters subcellular carbohydrate allocation. Previous work has established metabolite compartmentation as a crucial regulatory layer controlling metabolic flexibility during environmental acclimation (Knaupp et al., 2011; Hoermiller et al., 2017). Consistent with this concept, plastidial sucrose and vacuolar hexose pools exhibited opposing responses between single-grown and bulk-grown plants despite comparatively similar total hexose dynamics. These findings indicate that identical cellular metabolite concentrations may arise from fundamentally different intracellular allocation strategies. Particularly striking was the reciprocal regulation of plastidial and cytosolic sucrose pools, suggesting that intracellular carbon partitioning constitutes an actively regulated component of acclimation rather than a passive consequence of changing metabolite abundances. Such observations emphasize that analyses restricted to whole-tissue metabolite contents may overlook important regulatory differences operating at the compartment-specific level.

The output of the correlation analysis further supports this interpretation by revealing growth-type-dependent changes in metabolic coordination. In bulk-grown plants, enzyme activities showed substantially stronger associations with subcellular metabolite distributions than in isolated plants. Particularly the relationships between hexokinases, invertases and compartment-specific sugar pools indicated a tighter coupling between enzymatic capacity and intracellular metabolite partitioning. By contrast, weaker correlations in single-grown plants may indicate increased metabolic flexibility, allowing larger adjustments in metabolite pools despite comparable enzymatic capacities. While correlation analysis cannot establish causality, the observed reorganization of metabolic associations strongly suggests that the growth setup modifies the regulation of the subcellular sugar network. In this context, the data support the concept that acclimation should be considered as a systems property emerging from coordinated regulation across multiple metabolic processes rather than from changes in individual enzymes or membrane transporters (Fürtauer and Nägele, 2016; John et al., 2025).

The stronger accumulation of superoxide dismutase activity, malondialdehyde and anthocyanins in single-grown plants suggests that isolated plants experienced a greater requirement for photoprotective and antioxidant adjustment during cold exposure (see Figure 5). Importantly, these observations should not necessarily be interpreted as evidence of impaired acclimation. Instead, they are consistent with a more extensive physiological reprogramming required to establish a new metabolic equilibrium. Because photosynthetic parameters ultimately stabilized in both growth setups, the higher stress-marker accumulation in isolated plants likely reflects differences in acclimation trajectory rather than differences in acclimation success (Distelbarth et al., 2013; Herrmann et al., 2021).

The ecological implications of these findings deserve particular consideration. Most laboratory studies intentionally minimize environmental heterogeneity by cultivating individual plants under highly controlled conditions. Such approaches are indispensable for mechanistic studies because they reduce experimental variability and facilitate reproducibility. However, natural populations of *Arabidopsis thaliana* rarely experience such isolated growth conditions. Instead, neighbouring plants continuously modify light interception, resource availability, humidity, and developmental dynamics (Huber et al., 2021; Yan et al., 2023). Consequently, part of the phenotypic variation commonly attributed to genotype or environmental treatment may in fact arise from differences in the developmental and physiological context established by plant density and growth configuration. The present data therefore suggest that cultivation strategy should be regarded as an experimental factor influencing the expression of metabolic plasticity itself, particularly when studies aim to interpret natural variation or ecological adaptation.

Several limitations should nevertheless be considered. Bulk-grown and single-grown plants differed not only in spatial arrangement but also in developmental progression and sampling schedule, making it difficult to separate the effects of plant density, ontogeny and microenvironment. Likewise, the present study primarily quantifies metabolic states and enzymatic capacities rather than metabolic fluxes. Future studies combining isotope-based flux analysis with transcriptomics and quantitative systems modelling could therefore determine whether the observed differences in metabolite compartmentation directly arise from altered carbon fluxes or from changes in regulatory network architecture. Such approaches would also enable identification of the feedback mechanisms linking intracellular sugar partitioning, enzyme regulation and photosynthetic acclimation.

### Accession-specific information is predominantly encoded at the metabolic level

The classifier analysis revealed that the physiological and metabolic response to cold contained distinct signatures associated with both genetic background and growth setup. Importantly, however, the discriminatory information was not distributed uniformly across the measured variables. While photosynthetic and enzyme-related parameters provided comparatively little information for distinguishing the four *Arabidopsis thaliana* accessions, metabolic features - particularly organic acids and carbohydrates - showed substantially stronger accession-specific signatures. An even more pronounced pattern emerged for growth type, where organic acids and stress/acclimation markers almost completely separated plants grown under single and bulk conditions. The strongest discrimination among accessions was obtained from organic acids (macro-AUC = 0.841), followed by sugars (0.781) and subcellular sugar distribution (0.729). This observation is consistent with extensive evidence that cold acclimation involves profound reorganization of central carbon metabolism (Klotke et al., 2004; Saunders et al., 2022), and that the extent and dynamics of this reorganization vary among natural Arabidopsis accessions (Verslues and Juenger, 2011). Previous comparisons of accessions with different cold-acclimation capacities demonstrated accession-specific regulation of soluble carbohydrate metabolism, including differences in sucrose synthesis and sucrose/hexose interconversion during exposure to low temperature (Nägele and Heyer, 2013). More recently, metabolomic analysis of 241 natural *Arabidopsis* accessions demonstrated substantial genetically determined variation in metabolic plasticity at low temperature, with fumarate and sugar metabolism among the metabolic processes showing particularly pronounced accession-dependent responses (Weiszmann et al., 2023). The high discriminatory power of organic acids and sugars observed here is therefore consistent with central carbon metabolism representing an important component of natural variation in the response to low temperature.

Beyond, organic acids showed particularly strong discrimination of accessions with AUCs of >0.95. Organic acids are closely connected to respiratory carbon metabolism, cellular energy status and the provision of carbon skeletons for numerous biosynthetic processes (Medeiros et al., 2021). Cold acclimation results in broad changes in primary metabolism, including accumulation of several organic acids and intermediates associated with the tricarboxylic acid cycle. The strong accession-specific signature detected here may therefore reflect differences in the metabolic adjustment required to maintain carbon and energy homeostasis at low temperature. This interpretation is particularly interesting in the context of the reported association between biomass accumulation, fumarate metabolism and a naturally occurring promoter polymorphism of the *FUMARASE2* gene which encodes a cytosolic fumarase in *Arabidopsis* (Riewe et al., 2016). While the identification of the physiological role of FUM2 remains difficult, for example because it might also comprise compartment-specific redox regulation networks (Igamberdiev and Eprintsev, 2016), it might be of interest for future studies to analyse mutants under different growth conditions ranging from single to dense bulk conditions. The exceptionally high discriminatory performance of organic acids is particularly noteworthy because this feature group also provided the strongest growth type discrimination. Central organic acid metabolism therefore appears sensitive to both genetic background and experimental growth configuration. Growth-dependent differences could arise from variation in source–sink relationships, plant density, resource competition, microenvironmental conditions or developmental state, all of which can influence carbon demand and respiratory metabolism. More generally, cold acclimation requires a new balance between photosynthetic carbon acquisition, metabolic conversion, storage and growth because low temperature affects these processes to different extents (Seydel et al., 2022). Hence, the organic acid signature observed here may consequently integrate several physiological consequences of the growth setup rather than reflect a single growth-dependent process.

Interestingly, in contrast to organic acids and carbohydrates, photosynthetic parameters showed only little discriminatory capacity among accessions (macro-AUC = 0.423). Low temperature directly affects photosynthetic carbon fixation and creates a requirement to coordinate photosynthesis with downstream carbon utilization (Herrmann et al., 2020). Photosynthetic capacity can itself acclimate substantially during prolonged exposure to low temperature. Moreover, chlorophyll fluorescence parameters have previously been used to distinguish cold-sensitive and cold-tolerant *Arabidopsis* accessions under particular experimental conditions (Mishra et al., 2011). One possible interpretation for the low AUC observed here is that adjustment of photosynthetic function represents a comparatively conserved component of the cold response across the investigated accessions, whereas downstream carbon allocation and metabolic homeostasis are more genotype dependent.

### Implications and limitations of the classifier analysis

An unexpected observation was that classifiers incorporating all measured variables performed considerably worse than classifiers restricted to selected metabolic feature groups for both classification tasks. For accession classification, the all-feature model achieved a macro-AUC of only 0.485 compared with 0.841 for organic acids; for growth type, the corresponding values were 0.620 and 1.000. Thus, adding more physiological information did not improve classification. This may indicate that highly informative metabolic signals were diluted by variables containing little class-specific information. In a neural-network model with a relatively limited number of observations, increasing input dimensionality may additionally increase model complexity and susceptibility to overfitting (Hawkins, 2003). The superior performance of restricted feature sets should therefore not be interpreted as evidence that other physiological processes are biologically irrelevant. Instead, organic acids and sugars appear to provide a more concentrated discriminatory signal within the present dataset. Further, the very high AUCs obtained for growth-type classification, particularly the AUC of 1.000 for organic acids, also warrant caution. The independent test set comprised only 15% of the complete dataset, and perfect separation within a relatively small test subset does not necessarily imply perfect generalization to independent experiments. Variation in random train–validation–test partitioning could substantially affect performance estimates. Accordingly, these findings should be interpreted as evidence for strong candidate signatures rather than definitive biomarkers which might motivate future research in this context.

Overall, the classifier analysis demonstrates that cold-induced physiological and metabolic dynamics contain both conserved and context-dependent components. Photosynthetic responses showed comparatively little information about accession or growth setup, suggesting a substantial degree of conservation across the investigated conditions. In contrast, central carbon metabolism, particularly metabolism of organic acids and soluble sugars, retained pronounced accession-specific signatures, consistent with natural variation in metabolic cold acclimation reported previously. At the same time, the strong discrimination between single and bulk growth based on organic acids and stress/acclimation markers demonstrates that metabolic acclimation is highly sensitive to the experimental growth environment. These findings therefore emphasize that conservation of the physiological cold response does not necessarily imply conservation of the underlying metabolic state. Rather, different genotypes and growth configurations may achieve acclimation through distinct metabolic configurations, with organic acid and carbohydrate metabolism emerging as particularly informative indicators of this variation.

## Acknowledgements

We thank the members of Plant Evolutionary Cell Biology at LMU München and the members of AG Weckwerth at University of Vienna for fruitful discussions. Further, we thank the Graduate School Life Science Munich (LSM) for support. This work was funded by Deutsche Forschungsgemeinschaft (DFG), NA1545-5/1, and by the Austrian Science Fund (FWF), I 5234.

## Data availability statement

Data of this study is available in the supplements.

## Author contributions

**VB**: Performed experiments, statistics, data evaluation and wrote the manuscript. **WW:** outlined and supervised data evaluation and wrote the manuscript. **TN**: conceived the study, performed statistics and data evaluation, and wrote the manuscript.

## Supplementary Information

**Supplementary Figure SF1.**
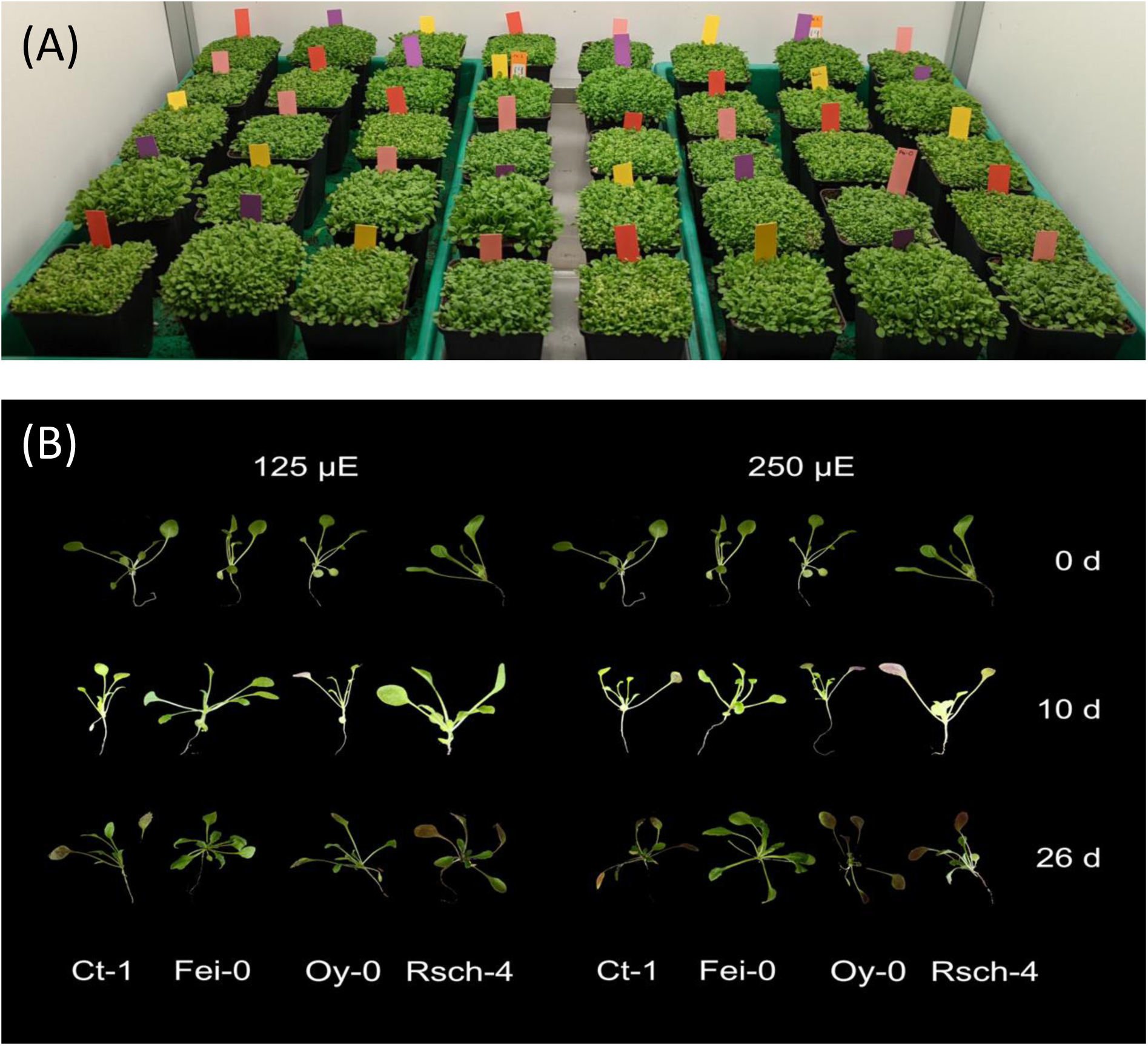
Phenotypes of *Arabidopsis* thaliana seedlings. **(A)** bulk grown plants. The size of the pots was 9×9 cm. **(B)** Individual plant phenotypes: Ct-1, Fei-0, Oy-0, Rsch-4 after 0d, 10d, 26d in the cold under 125 µE /250 µE growth light intensity.

**Supplementary Figure SF2.**
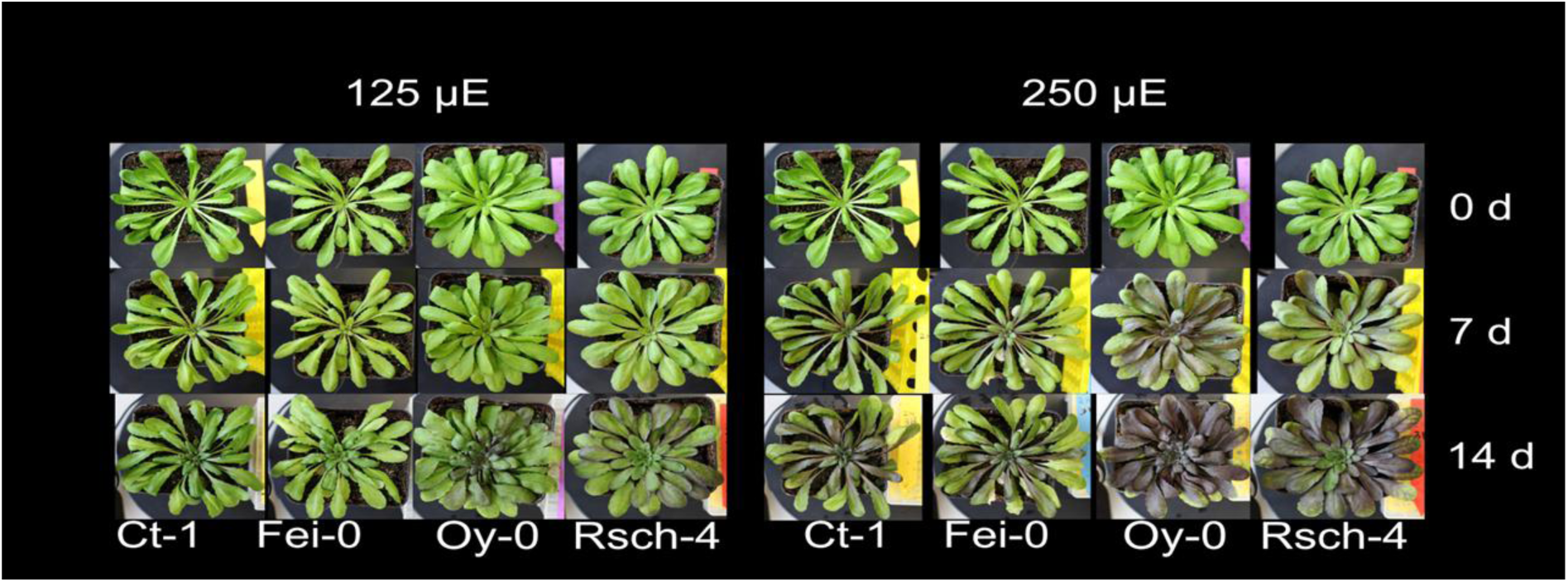
Phenotypes of mature plants of *Arabidopsis* thaliana. Single plant phenotypes: photos were taken of accessions: Ct-1, Fei-0, Oy-0, Rsch-4 after 0d, 7d, 14d in the cold under 125 µE /250 µE growth light intensity. The size of the pots was 9×9 cm.

**Supplementary Table ST1. Statistical analysis of physiological and metabolic variables using analysis of variance (ANOVA).** For each variable, the table reports the tested main effects and interaction terms involving accession, growth type, time point (tp), and experimental condition, together with the corresponding p-values and partial eta-squared (η²ₚ) effect sizes. Variables are grouped according to their respective physiological or metabolic categories. Statistical significance is indicated as p < 0.05 (), p < 0.01 (), and p < 0.001 (); non-significant effects are shown without a symbol. Partial η² values provide an estimate of effect size, with values of approximately 0.01, 0.06, and 0.14 corresponding to small, medium, and large effects, respectively.

**Supplementary Table ST2. Complete dataset of physiological and metabolic measurements used in the study.** The table contains the individual measurements for all Arabidopsis accessions, growth types, time points, experimental conditions, and biological replicates included in the analyses. Measured variables comprise carbohydrate and organic acid contents, enzyme activities, stress and acclimation markers, biomass parameters, photosynthetic and chlorophyll fluorescence parameters, CO₂ assimilation (nps), and subcellular sugar distributions.

**Supplementary Table ST3. Classifier performance for discrimination between *Arabidopsis* accessions using different physiological and metabolic feature groups.** Classification performance is reported for classifiers trained using all measured features or individual feature groups, including photosynthetic parameters, enzyme activities, sugars, organic acids, stress and acclimation markers, and subcellular sugar distribution. Class-specific areas under the receiver operating characteristic curve (AUCs) are provided for the accessions Ct-1, Fei-0, Oy-0, and Rsch-4. The macro-AUC represents the unweighted mean of the four accession-specific AUC values. Overall classification performance is additionally summarized by accuracy, macro-precision, macro-recall, and macro-F1 score. All performance metrics were calculated using the independent test dataset. AUC values of 0.5 indicate discrimination at chance level, whereas values approaching 1.0 indicate increasing discriminatory performance.

**Supplementary Table ST4. Classifier performance for discrimination between single and bulk growth types using different physiological and metabolic feature groups.** Classification performance is reported for classifiers based on all measured features or individual feature groups, including photosynthetic parameters, enzyme activities, sugars, organic acids, stress and acclimation markers, and subcellular sugar distribution. Class-specific areas under the receiver operating characteristic curve (AUCs) are provided for single and bulk growth. The macro-AUC represents the unweighted mean of the two class-specific AUC values. Overall classifier performance is additionally summarized by accuracy, macro-precision, macro-recall, and macro-F1 score. All performance metrics were calculated using the independent test dataset. AUC values of 0.5 indicate discrimination at chance level, whereas values approaching 1.0 indicate increasing discriminatory performance.

